# Dynamic Reassociation of the Nuclear Lamina with Newly Replicated DNA

**DOI:** 10.1101/2022.11.11.516151

**Authors:** Callie M. Lovejoy, Prabakaran Nagarajan, Mark R. Parthun

## Abstract

The physical association of specific regions of chromatin with components of the nuclear lamina provides the framework for the 3-dimensionl architecture of the genome. The regulation of these interactions plays a critical role in the maintenance of gene expression patterns and cell identity. The breakdown and reassembly of the nuclear membrane as cells transit mitosis plays a central role in the regulation of the interactions between the genome and the nuclear lamina. However, other nuclear processes, such as transcription, have emerged as regulators of the association of DNA with the nuclear lamina. To determine whether DNA replication also has the potential to regulate DNA-nuclear lamina interactions, we adapted proximity ligation-based chromatin assembly assays to analyze the dynamics of nuclear lamina association with newly replicated DNA. We observe that lamin A/C and lamin B, as well as inner nuclear membrane proteins LBR and emerin, are found in proximity to newly replicated DNA. While core histones rapidly reassociate with DNA following passage of the replication fork, the complete reassociation of nuclear lamina components with newly replicated DNA occurs over a period of approximately 30 minutes. We propose models to describe the disassembly and reassembly of nascent chromatin with the nuclear lamina.

## INTRODUCTION

Eukaryotic genomes are organized in a non-random fashion in the three-dimensional environment of the nucleus. At the most fundamental level, the genome is partitioned into two compartments, A and B. The A compartment is euchromatic and localized in the interior of the nucleus, while the B compartment is heterochromatic and localized to the nuclear periphery. This genome architecture is functionally important and plays a critical role in regulating gene expression programs and maintaining genome integrity(1,2).

A key driving force controlling this three-dimensional structure is the tethering of specific regions of the genome to the surface of the inner nuclear membrane through interactions with the nuclear lamina. The nuclear lamina is a meshwork of intermediate filaments composed of A- and B-type nuclear lamins. The lamins are anchored to the nuclear periphery by interactions with inner nuclear membrane proteins, including the lamin B receptor (LBR), emerin, Lap2β, Man1, and the LINC complex(3-5). The lamins also interact with specific regions of the genome termed lamin-associated domains (LADs)(6). LADs range in size from ∼0.1 to 10 Mb and encompass 30% to 40% of the mammalian genome(7,8).

Regions of the genome associated with the nuclear lamina are typically heterochromatic and associated with the B compartment. LADs have a low gene density and most genes in LADs are poorly expressed. LADs are typically late replicating regions of the genome. LAD chromatin is enriched for the repressive histone modifications H3 K9me2, H3 K9me3, and H3 K27me3(3,8).

Interactions with the nuclear lamina involve aspects of both the DNA sequence and chromatin structure of LADS. LAD DNA tends to have a high A-T content and a number of specific DNA sequences have been isolated that are targeted to the nuclear lamina(8-12). However, the primary determinant of LAD association with the nuclear lamina appears to be chromatin state(11). The di- and tri-methylation of H3 K9, a mark of constitutive heterochromatin, is required for localization of LADs to the nuclear periphery. The association of constitutive heterochromatin with the nuclear lamina is mediated by the H3 K9me2/3 reader protein HP1, which can directly interact with the lamina-associated proteins LBR and PRR14. There are also direct contacts between histones and nuclear lamina components as LBR can also bind to histone H4 di-methylated on K20, another modification enriched in constitutive heterochromatin(10,11,13,14).

The interactions between chromatin and the nuclear lamina are highly dynamic(5). Some regions of the genome are found associated with the nuclear lamina in most cell types and are known as constitutive LADs (cLADs). Other regions of the genome, facultative LADs (fLADs), are only associated with the nuclear lamina in certain cell types or at specific points during development(7,15). These alterations in genome architecture are likely to play an important role in the specification and maintenance of cell identity.

Nuclear lamina-chromatin interactions are also dynamic with respect to the cell cycle. The most dramatic changes in chromatin-nuclear lamina interactions occur during mitosis. As cells enter mitosis and chromosomes condense, interactions with the nuclear lamina and inner nuclear membrane are lost as the nuclear envelope breaks down and the lamins are dispersed into the cytoplasm. As cells prepare to exit mitosis, the nuclear envelope reforms in the daughter cells and interactions between LADs and the nuclear lamina are reestablished(5,8). The organization of LADs can change when cells pass through mitosis as tracking of LADs in single cells indicate that many genomic regions localized to the nuclear periphery in mother cells become localized to the interior following cell division(14,16).

Chromatin-nuclear lamina interactions are also dynamic outside of mitosis. While many studies have demonstrated that the association of loci with the nuclear lamina leads to a down-regulation of transcription, recent studies have shown that transcription can also directly regulate the association of genes with the nuclear lamina(17-20). Targeting a strong transcriptional activator to several loci caused a decrease in the association of the targeted genes and nearby flanking sequences with nuclear lamina components. Conversely, repressing active genes led to an increase in nuclear lamina interactions(21). Hence, while the underlying mechanism is not known, the process of transcribing a gene can modulate localized interactions of chromatin with the nuclear lamina.

While much attention has been focused on the function of the nuclear lamina in the regulation of transcription, how the nuclear lamina influences DNA replication is poorly understood. The observation that LADs replicate late in S-phase suggests that the association of chromatin with the nuclear lamina creates an environment that is repressive for the initiation of DNA replication(6,15,22). Nuclear lamina components are required for genome integrity and recent results indicate that lamin A/C can interact with RPA and Rad51 to promote the stability of stalled replication forks(2,23).

An interesting open question is whether the process of DNA replication regulates chromatin-nuclear lamina interactions. During progression of a replication fork, parental nucleosomes are displaced as the CMG helicase unwinds the double stranded DNA. The released histones dissociate into histone H3/H4 tetramers and H2A/H2B dimers. The parental H3/H4 tetramers, which possess the bulk of the histone post-translational modifications required to epigenetically specify heterochromatin structure, are captured by components of the replisome that possess histone chaperone activity and redeposited on the newly replicated DNA behind the replication fork(24-31).

Nucleosome density on the two daughter duplexes is maintained by the deposition of an equal quantity of newly synthesized H3/H4 tetramers by the CAF-1 chromatin assembly complex(29,30). The impact of chromatin disassembly and reassembly on the interaction of heterochromatin with the nuclear lamina is not known.

To begin to address this question, we have adapted proximity ligation-based chromatin assembly assays to analyze the dynamic association of nuclear lamina components with DNA following passage of a replication fork(32-35). We observe that the levels of these nuclear lamina components associated with newly synthesized DNA significantly increases in the first 30 minutes following replication and then plateaus. This pattern is distinct from that observed for the association of core histones with newly replicated DNA. We propose several models to describe the association of newly synthesized DNA with the nuclear lamina following DNA replication.

## RESULTS

The maintenance of nuclear architecture following DNA replication is necessary to preserve the gene expression patterns required for cellular identity. Reproducing the three-dimensional structure of the genome following DNA replication must involve the accurate restoration of chromatin structures that are competent for making the appropriate contacts with the nuclear periphery. The mechanisms that coordinate the assembly of nascent chromatin structure on newly replicated DNA with reattachment to the nuclear lamina are unknown.

### Nuclear lamins are in close proximity to newly replicated DNA

The association of nuclear lamins with newly replicated DNA was first suggested by the immunofluorescent co-localization of lamin B1 with BrdU labeled DNA in mid to late S phase cells(36). More recent studies of nascent chromatin proteomics have used iPOND (isolation of <u>p</u>roteins <u>o</u>n <u>n</u>ascent <u>D</u>NA) or NCC (<u>n</u>ascent <u>c</u>hromatin <u>c</u>apture) to detect the association of lamin A/C and lamin B with newly replicated DNA(23,37,38).

To study the association of nuclear lamins with newly replicated DNA, we adapted the proximity ligation-based chromatin assembly assay (PL-CAA) (32,34,39,40). The proximity ligation technique determines whether two molecules reside close to each other in the cell by employing two species-specific secondary antibodies that are fused to oligonucleotides. If the secondary antibodies recognize primary antibodies that are in close proximity, the oligonucleotides can both bind to a nicked circular DNA, creating a template for rolling circle replication. This amplifies sequences that can be bound by a fluorescent probe and visualized. To use this as a chromatin assembly assay, newly replicated DNA is labeled by incorporation of the thymidine analog IdU. The proximity of proteins to newly replicated DNA is detected using antibodies against the protein of interest and antibodies recognizing IdU. PL-CAA has important advantages over other nascent chromatin proteomics techniques. The PL-CAA does not involve the Click chemistry used in iPOND or NCC, which can mask epitopes on a high fraction of proteins. In addition, PL-CAA does not require large numbers of cells. Importantly, PL-CAA offers single cell resolution and provides information about the sub-cellular localization of interactions.

We incubated mouse embryonic fibroblasts (MEFs) with IdU for 30 minutes to label newly replicated DNA and performed PLA-CAA using antibodies recognizing either lamin A/C or Lamin B1 and IdU (Figure 1A). We observed abundant PLA-CAA signal with both nuclear lamins, specifically in IdU positive cells (Figure 1B and 1C) Importantly, this signal was enriched at the nuclear periphery. These results confirm the proximity of lamin A/C and lamin B1 with newly replicated DNA in MEFs and demonstrate that PL-CAA can be used to evaluate the dynamics of nuclear lamin association with DNA following the passage of a replication fork(23,36,38).

**Figure 1.**
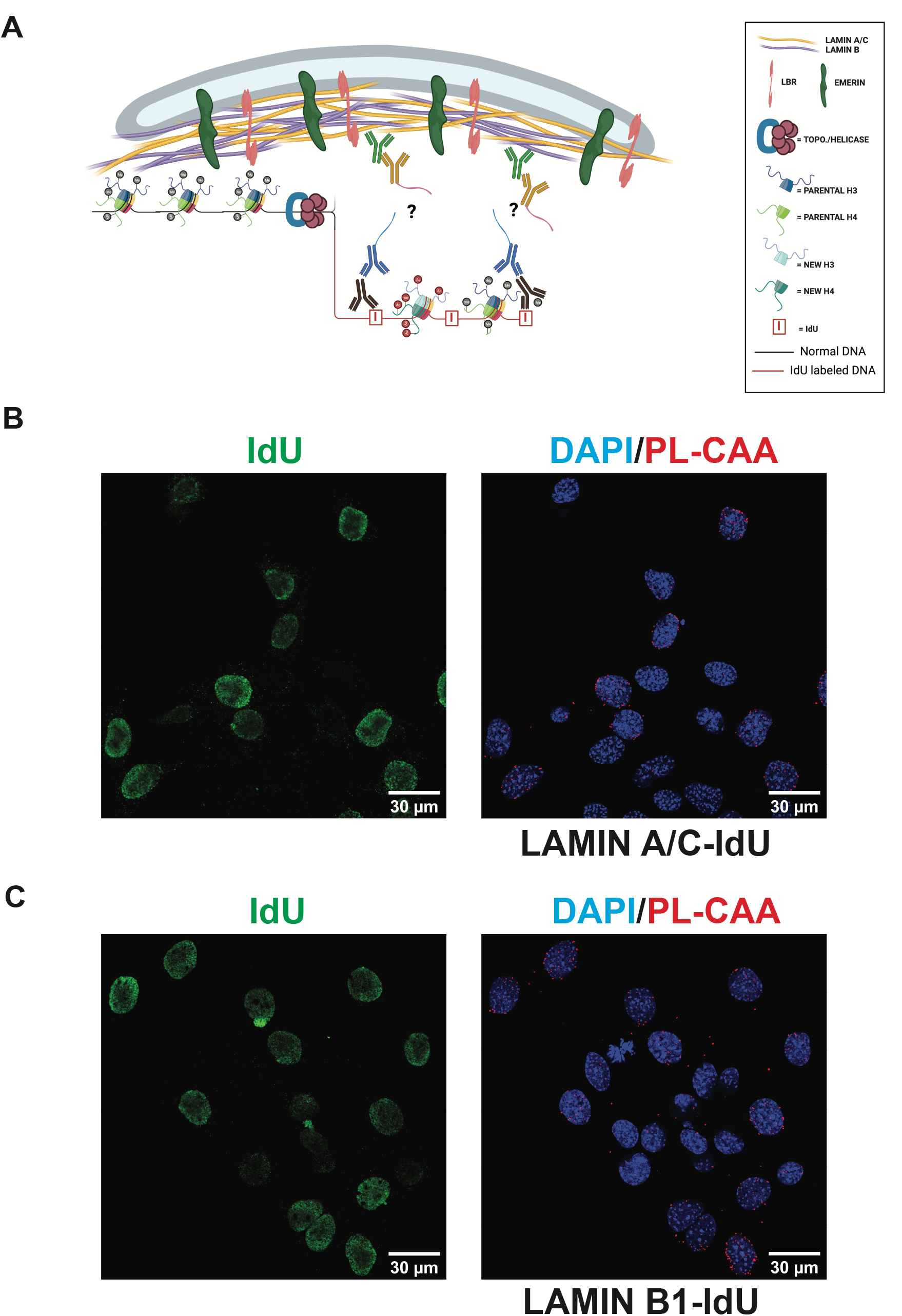
(A) Model representation of antibody detection of IdU and proteins of interest using the PL-CAA method. (B) Representative images of PL-CAA in MEFs detecting close proximity of Lamin A/C to newly replicated DNA, indicated by red foci. PL-CAA foci are specific to IdU positive cells (green). (C) Representative images of PL-CAA in MEFs detecting close proximity of Lamin B1 to newly replicated DNA, indicated by red foci. PL-CAA foci are specific to IdU positive cells (green).

### Reassociation of nuclear lamins with newly replicated DNA

To analyze the dynamics of protein association with newly replicated DNA, we used a pulse-chase strategy, where newly replicated DNA was labeled by incubation of MEFs in media containing IdU for 30 minutes. IdU-containing media was then washed out and cells were then incubated in media containing thymidine for up to 2 hours allowing us to monitor nascent chromatin maturation.

As a positive control to demonstrate the use of PL-CAA to monitor the dynamic association of proteins with DNA post-replication, we analyzed a core histone. In the wake of the replisome, both the leading and lagging strands are rapidly repackaged into nucleosomes through the recycling of parental histones and the deposition of newly synthesized histones. This repackaging is rapid and highly efficient, preventing the accumulation of any appreciable stretches of unchromatinized double strand DNA(29,30,41). This was reflected in PL-CAAs using antibodies to histone H3 and IdU (Figure 2A). There is a high level of interaction between histone H3 and newly replicated DNA immediately following the IdU pulse and this level remained constant over the 2 hour thymidine chase.

**Figure 2.**
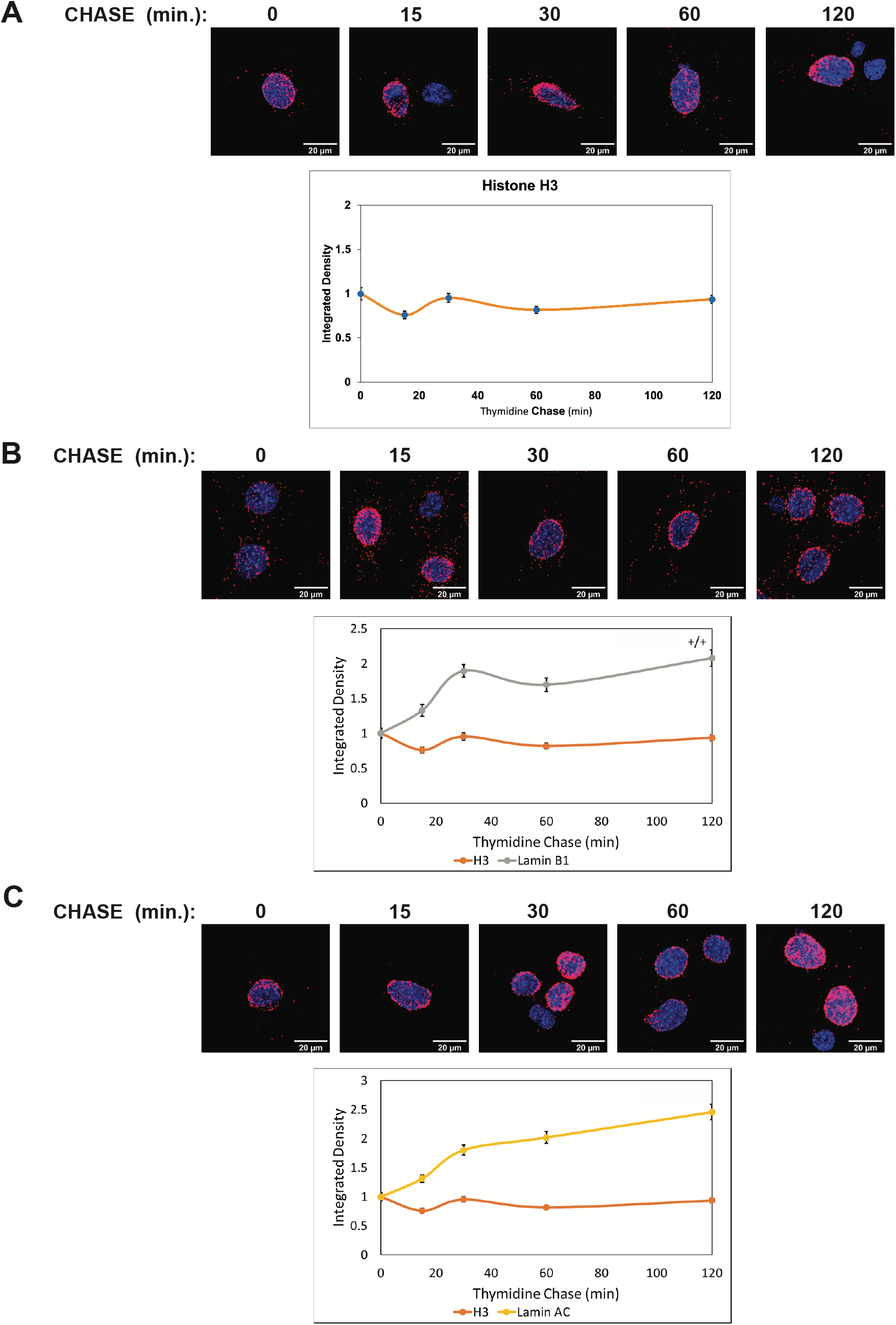
(A) Representative images of Histone H3 PL-CAA at 0 min, 15 min, 30 min, 60 min, and 120 min after IdU labelling as a control for immediate reassembly of nascent DNA with the protein of interest. Quantification of integrated density of Histone H3 PL-CAAs at 0-120 minutes post-DNA label. (B) Representative images of Lamin B1 PL-CAA at 0 min, 15 min, 30 min, 60 min, and 120 min after IdU labelling Quantification of integrated density of Lamin B1 PL-CAAs at 0-120 minutes post-DNA label. (C) Representative images of Lamin A/C PL-CAA at 0 min, 15 min, 30 min, 60 min, and 120 min after IdU labelling Quantification of integrated density of Lamin A/C PL-CAAs at 0-120 minutes post-DNA label. (n > 168 per timepoint within a PL-CAA reaction)

The kinetics of nuclear lamin reassociation with newly replicated DNA differed markedly from that observed for histone H3 (Figure 2B and 2C). There was clearly detectable PL-CAA signal for both Lamin A/C (Figure 2C) and Lamin B1 (Figure 2B) immediately after the IdU pulse. However, the PL-CAA signal doubled during the first 30 minutes of nascent chromatin maturation and then plateaued. These results suggest that newly synthesized DNA-nuclear lamina interactions are not restored immediately after passage of the replication as seen for histone deposition. This suggests that passage of a replication fork provides a window of opportunity for regulating the association of genomic DNA with the nuclear periphery.

### Inner nuclear membrane proteins are in proximity to newly replicated DNA

In addition to lamin A/C and lamin B, proteomic analyses of nascent chromatin also identified inner nuclear membrane proteins as physically associated with newly replicated DNA, including LBR and emerin. To determine whether these interactions could be visualized by PL-CAA, we incubated MEFs for 30 minutes with IdU and performed proximity ligation assays using antibodies against IdU and either LBR or emerin. PLA-CAA signals that were specific for IdU positive cells were observed for both LBR and emerin, confirming that these inner nuclear membrane proteins come into association with genomic DNA following replication(Figure 3A and 3B).

**Figure 3.**
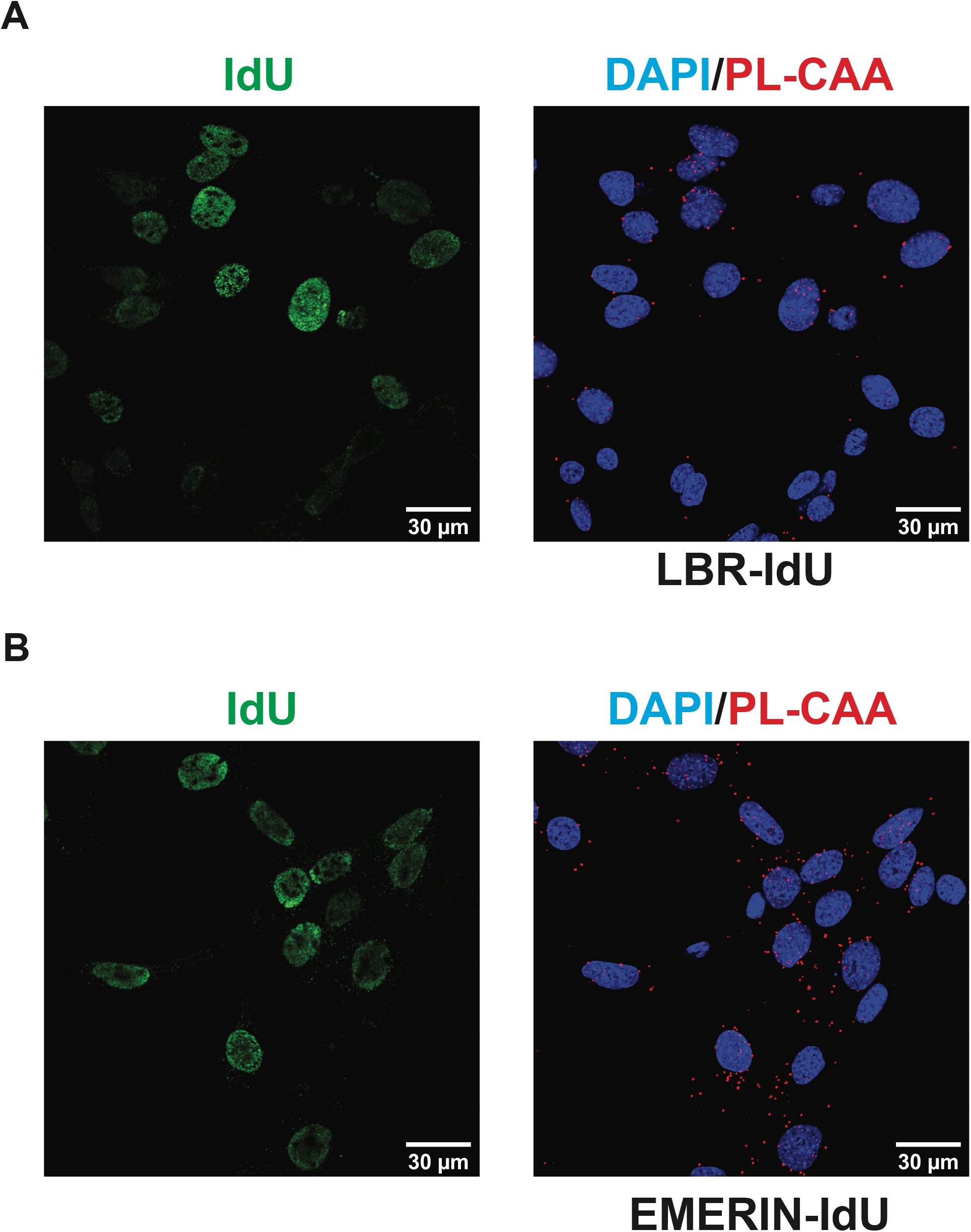
(A) Representative images of PL-CAA in MEFs detecting close proximity of LBR to newly replicated DNA, indicated by red foci. PL-CAA foci are specific to IdU positive cells (green). (B) Representative images of PL-CAA in MEFs detecting close proximity of Emerin to newly replicated DNA, indicated by red foci. PL-CAA foci are specific to IdU positive cells (green).

### Reassociation of inner nuclear membrane proteins with newly replicated DNA

We used the same pulse-chase strategy to determine the dynamics of LBR and emerin association with newly synthesized DNA during nascent chromatin maturation. The association of LBR with replicated DNA increased approximately 3-fold over the first 30 minutes of the thymidine chase and then remained stable (Figure 4A). The pattern was somewhat different for emerin. While the association of emerin with IdU-labeled DNA increased during nascent chromatin maturation, there was a lag period of 15 minutes before the emerin PL-CAA signal increased (Figure 4B). After 2 hours of nascent chromatin maturation, the emerin PL-CAA signal had increased approximately 4-fold. These results demonstrate that non-lamin components of the nuclear lamina also dynamically associate with newly replicated DNA and that specific nuclear lamina components have distinct patterns of reassociation with the genome following passage of a replication fork.

**Figure 4.**
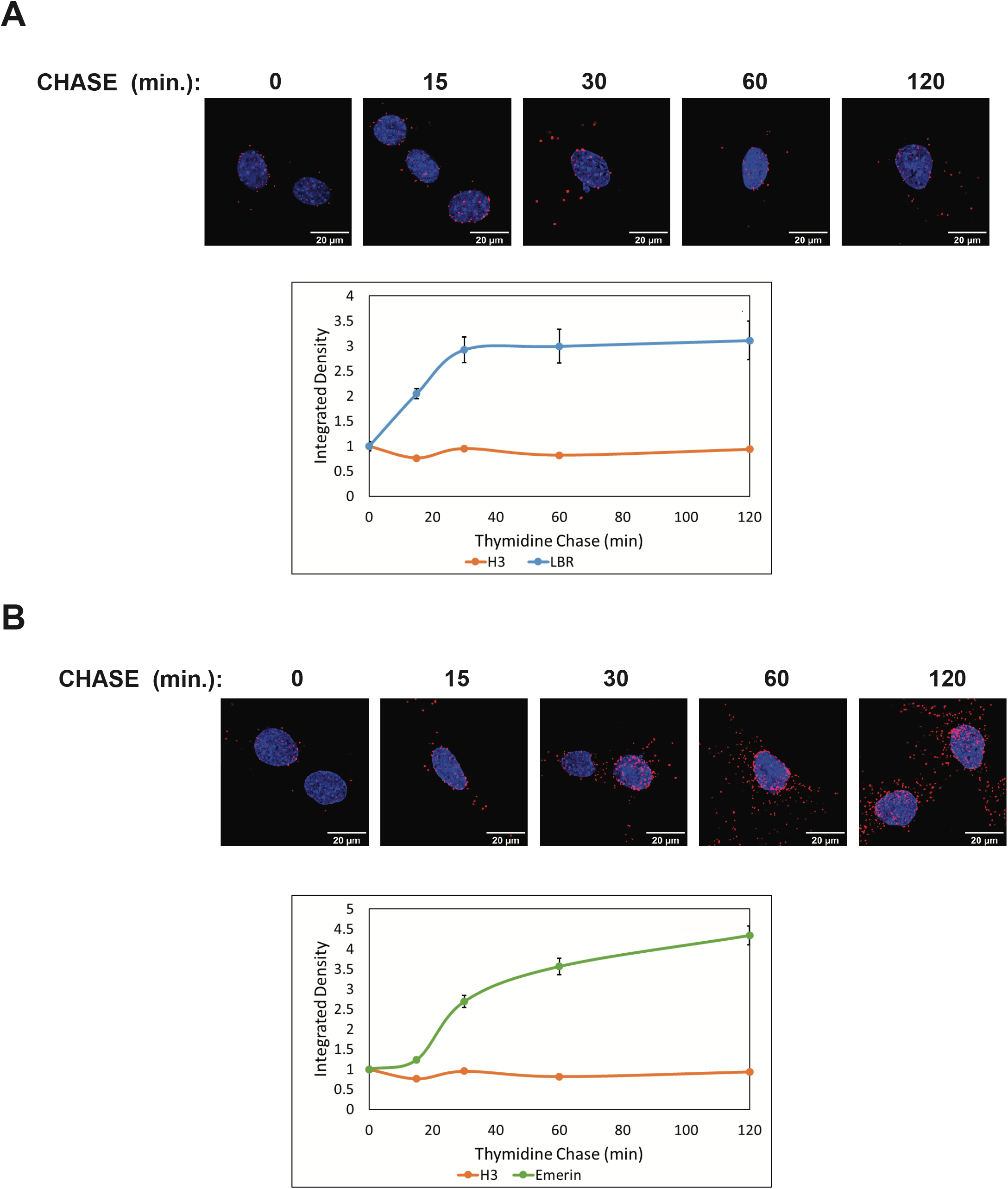
(A) Representative images of LBR PL-CAA at 0 min, 15 min, 30 min, 60 min, and 120 min after IdU labelling Quantification of integrated density of Lamin B1 PL-CAAs at 0-120 minutes post-DNA label. (B) Representative images of Lamin A/C PL-CAA at 0 min, 15 min, 30 min, 60 min, and 120 min after IdU labelling Quantification of integrated density of Emerin PL-CAAs at 0-120 minutes post-DNA label. (n > 167 per timepoint within a PL-CAA reaction)

## DISCUSSION

The fundamental question of whether passage of a replication fork disrupts the association of chromatin with the nuclear lamina has not been explored. We used proximity ligation-based assays to demonstrate that DNA replication alters the interaction between components of the nuclear lamina and newly replicated DNA and that there is a lag period before the interaction is restored, as the reestablishment of the nuclear lamina-chromatin interaction is not completed until approximately 30 minutes after replication.

The delayed reassociation of chromatin with the nuclear lamina following DNA replication provides an opportunity for the regulation of 3-dimensional genome archit4ecture. This is consistent with results of an siRNA screen that identified factors required for 3-D gene positioning. Several factors involved in DNA replication and replication-coupled chromatin assembly, such as PCNA, CHAF1A and ASF1A, were influence gene positioning. Importantly, DNA replication itself was identified as a critical determinant of 3-D genome architecture(42).

We propose a model to describe the effect of DNA replication on the association of chromatin with the nuclear lamina (Figure 5). As the association of heterochromatic LADs with the nuclear periphery is driven by physical interactions between protein components of the nuclear lamina and proteins associated with chromatin, the disassembly of nucleosomes resulting from passage of a replication temporarily displaces newly replicated DNA from the nuclear lamina (Figure 5A, top). Following a 30 minute pulse with IdU, a significant PL-CAA signal is detectable for both the lamin A/C-IdU and lamin B1-IdU interactions, suggesting partial restoration of the nuclear lamina-nascent chromatin interaction during the 30 minute IdU pulse. However, unlike the rapid reassembly of histones on newly replicated DNA, the PL-CAA signal increases during the subsequent thymidine chaser, indicating that the nuclear lamina-nascent chromatin interaction reassembles with slower kinetics that nucleosome assembly.

**Figure 5.**
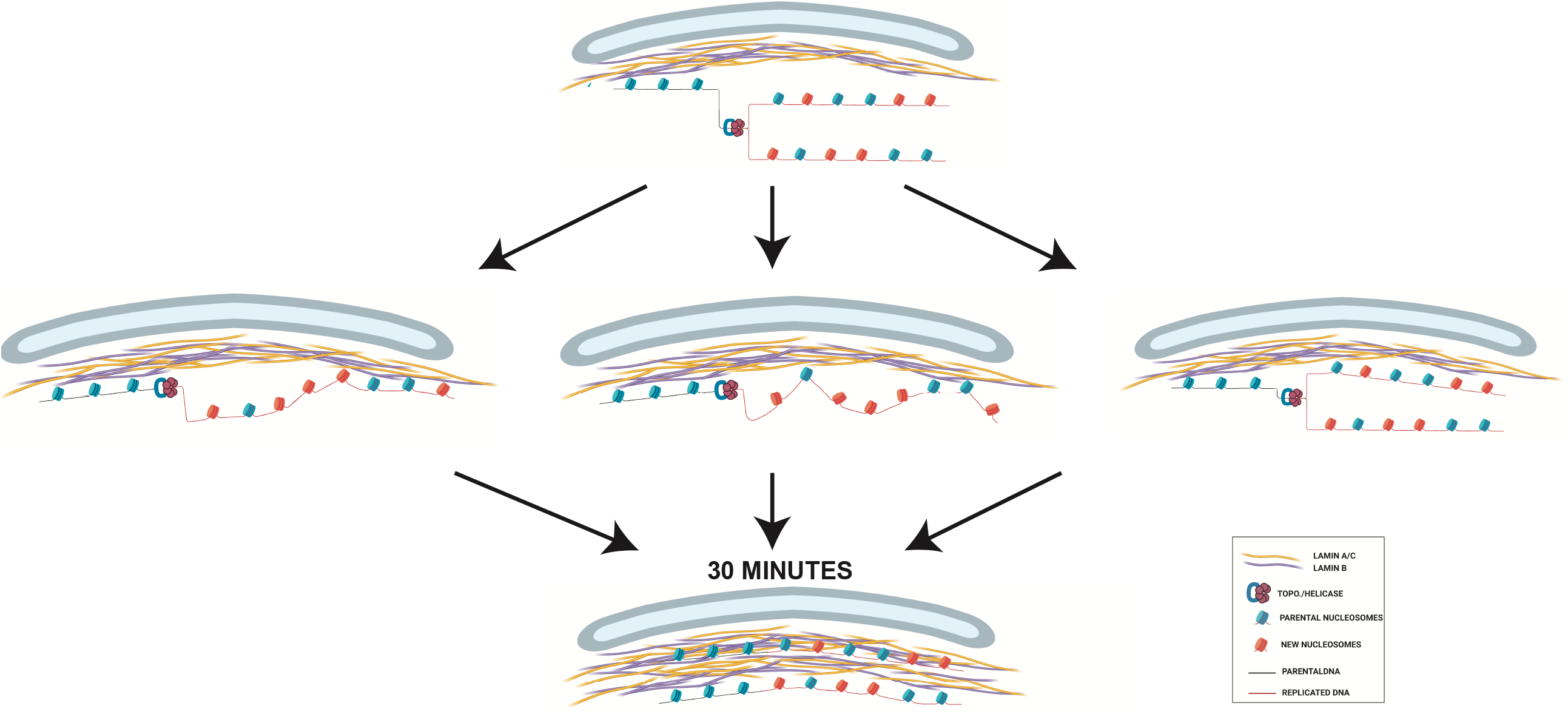
(A) Proposed model of chromatin reassociation with the nuclear lamina post-DNA replication.

The 2-fold increase in the PL-CAA signal for both lamin A/C-IdU and lamin B1-IdU in the first 30 minutes following the IdU pulse suggests that only half of the nuclear lamina-nascent chromatin interaction was restored during the period of the IdU pulse and that half of the interaction was restored during the first 30 minutes of the thymidine chase. There are three scenarios that are consistent with these observations. First, after disruption by the replication fork, the chromatin could reassociate directionally, with the earliest replicating regions binding to the nuclear lamina first. This “zippering up” of the interactions between nucleosomes and the nuclear lamina would occur at a relatively slow rate, resulting in a lag between the replication-coupled assembly of the replicated DNA into nucleosomes and their binding to the nuclear lamina (Figure 5, middle left).

Nascent chromatin is composed of a 1:1 mixture of parental histones and newly synthesized histones(29,30). Therefore, another potential scenario is that parental histones rapidly reassociate with the nuclear lamina, while the newly synthesized histones require a maturation period before they are competent for binding to the nuclear lamina (Figure 5, middle center). Following their deposition onto DNA, the modification state of newly synthesized histones is highly dynamic. New histones H3 and H4 acquire specific pre-deposition patterns of acetylation that are removed over a period of roughly 1 hour following chromatin assembly(43-45). In regions of constitutive heterochromatin, the newly synthesized histones must also acquire methylation on H3 K9, which is likely to be a prerequisite for association with the nuclear lamina(8). The acetylation state of newly synthesized histones may influence their subsequent methylation as HAT1 and the acetylation of newly synthesized H3 and H4 regulates H3 K9me2/3 in large chromatin domains, termed HADs (<u>H</u>AT1-dependent <u>A</u>ccessibility <u>D</u>omains), that show significant overlap with LADs(46). Additionally, HDAC2 and HDAC3 are physically associate with the nuclear lamina and may play a role in converting acetylated newly synthesized histones into a form capable of interacting with the nuclear lamina(47,48).

A third possibility is that nascent chromatin on the leading and lagging strands reassociate with the nuclear lamina with different kinetics, with one duplex rapidly rebinding the nuclear lamina and the other strand rebinding slowly (Figure 5, middle right). Distinct pathways are responsible for the recycling of parental histones onto the leading and lagging strands. A pathway involving MCM2 and DNA Polα directs parental histones to the lagging stand while a pathway involving Polε3 and polε4 targets parental histones to the leading strand(28,29,49,50). This raises the possibility that there are also separate pathways that facilitate the reassociation of the leading and lagging strands with the nuclear lamina.

Following the IdU pulse, the PL-CAA signal for LBR-IdU and emerin-IdU increase by 3 to 4 fold. The very low level of PL-CAA signal immediately following the IdU pulse suggests that the difference in the magnitude of the increase between the lamins and the inner nuclear membrane proteins may be due to a more complete disruption of the connection between the newly replicated DNA and LBR and emerin. Importantly, the increase in PL-CAA signal for LBR and emerin largely occurs during the first 30 minutes of the thymidine chase, suggesting that this is the critical time frame for restoration of the interaction between nascent chromatin and the nuclear periphery.

Whether the disassembly and reassembly of nuclear lamina-chromatin interactions that occurs during DNA replication provides a mechanism for regulating patterns of genome localization to the nuclear periphery is an open question. However, this is consistent with the importance of the CAF-1 chromatin assembly complex in the maintenance of cell identity(51). In addition, recent results demonstrate that cell fate reprogramming can be regulated by the speed of DNA replication forks(52). Further studies are required to elucidate the role of DNA replication in the regulation of nuclear lamina-chromatin interactions.

## Experimental Procedures

### Cell Culture Conditions

Mouse embryonic fibroblasts were prepared as previously described (45). Cells were grown in Dulbecco’s modified Eagle’s medium (Sigma) supplemented with 10% fetal bovine serum (Sigma) and penicillin/streptomycin (Gibco).

### PL-CAA

Three independent MEF cell lines were seeded in equal quantities on coverslips and allowed to attach for 24 h. Cells were then incubated with 10 uM IdU (Sigma; catalog number I7125) for 30 min. For thymidine chases, IdU containing medium was replaced with fresh medium for the indicated times. Cells were then permeabilized with 0.5% Triton X-100 and fixed with 4% PFA simultaneously for 15 min, rinsed with PBS, and fixed again with 4% PFA for 10 min at room temperature. After several PBS washes, cells were incubated with 1 N HCl for 10 min, washed with PBS until pH neutralizes, and blocked with 5% BSA for 1 h at room temperature. BSA was removed with PBS washes and primary antibodies detecting IdU and a protein of interest were diluted in 1% BSA, 0.3% Triton X-100 and added to cells overnight at 4 °C. The following day, primary antibodies were removed with PBS and cells were subjected to the Duolink™ Proximity Ligation Assay protocol according to the manufacturer’s instructions (Sigma: DUO92008, DUO92004, DUO92002, and DUO82049). Details of the antibodies used in this study are found in Table 1. After amplification, cells were incubated with Alexa Fluor 488-conjugated anti-mouse secondary antibody (1:250, Molecular Probes) for 1 h at room temperature, antibody was removed with PBS, nuclei were stained with 20 mM Hoechst 33342 Fluorescent Stain and mounted on slides using Vectashield. Slides were analyzed under a Zeiss LSM 900 Airyscan 2 Point Scanning Confocal microscope. Images were acquired using Zen Blue 3.0 and quantification was completed using ImageJ version 1.52t according to a previously described protocol (40). Data was analyzed and plots generated using RStudio Version 1.2.5042 running R Version 4.0.0.

**Table 1:** Antibody Information for PL-CAA Experiments

| Antigen | Host Species | Company | Cat. # | Dilution |
| --- | --- | --- | --- | --- |
| Emerin | rabbit | ProteinTech | 10351-1-AP | 1/100 |
| Histone H3 | rabbit | abcam | ab1791 | 1/300 |
| IdU/BrdU | mouse | Beckton Dickerson | 347580 | 1/20 |
| PCNA | rabbit | Santa Cruz Biotechnology | sc-7907 | 1/50 |
| LaminAC | rabbit | abcam | ab133256 | 1/200 |
| LaminB1 | rabbit | abcam | ab65986 | 1/200 |
| LBR | rabbit | ProteinTech | 12398-1-AP | 1/100 |

## Acknowledgements

This work was supported by grant GM R01062970 (M.R.P.). Support for microscopy was provided by grants P30 NS104177 and S10 OD026842.

## Author Contributions

C.M.L. helped conceive of the experiments, performed the investigations, developed methodology, analyzed data, visualized data, and helped write the manuscript. P.N. provided resources and helped write the manuscript. MRP helped conceive of the experiments, supervised the experiments, and helped write the manuscripts.

